# Mapping global shifts in *Saccharomyces cerevisiae* gene expression across asynchronous time trajectories with diffusion maps

**DOI:** 10.1101/2021.02.11.430862

**Authors:** Taylor Reiter, Rachel Montpetit, Ron Runnebaum, C. Titus Brown, Ben Montpetit

## Abstract

Grapes grown in a particular geographic region often produce wines with consistent characteristics, suggesting there are site-specific factors driving recurrent fermentation outcomes. However, our understanding of the relationship between site-specific factors, microbial metabolism, and wine fermentation outcomes are not well understood. Here, we used differences in *Saccharomyces cerevisiae* gene expression as a biosensor for differences among Pinot noir fermentations from 15 vineyard sites. We profiled time series gene expression patterns of primary fermentations, but fermentations proceeded at different rates, making analyzes of these data with conventional differential expression tools difficult. This led us to develop a novel approach that combines diffusion mapping with continuous differential expression analysis. Using this method, we identified vineyard specific deviations in gene expression, including changes in gene expression correlated with the activity of the *non-Saccharomyces* yeast *Hanseniaspora uvarum*, as well as with initial nitrogen concentrations in grape musts. These results highlight novel relationships between site-specific variables and *Saccharomyces cerevisiae* gene expression that are linked to repeated wine fermentation outcomes. In addition, we demonstrate that our analysis approach can extract biologically relevant gene expression patterns in other contexts (e.g., hypoxic response of *Saccharomyces cerevisiae*), indicating that this approach offers a general method for investigating asynchronous time series gene expression data.

**Importance:** While it is generally accepted that foods, in particular wine, possess sensory characteristics associated with or derived from their place of origin, we lack knowledge of the biotic and abiotic factors central to this phenomenon. We have used *Saccharomyces cerevisiae* gene expression as a biosensor to capture differences in fermentations of Pinot noir grapes from 15 vineyards across two vintages. We find that gene expression by non-*Saccharomyces* yeasts and initial nitrogen content in the grape must correlates with differences in gene expression among fermentations from these vintages. These findings highlight important relationships between site-specific variables and gene expression that can be used to understand, or possibly modify, wine fermentation outcomes. Our work also provides a novel analysis method for investigating asynchronous gene expression data sets that is able to reveal both global shifts and subtle differences in gene expression due to varied cell – environment interactions.

## Introduction

During a wine fermentation, *Saccharomyces cerevisiae* metabolizes sugars and other nutrients to obtain energy and increase biomass, while also dealing with a common set of stresses caused by the must/wine environment. Given these general features of the system, the cellular activities of *S. cerevisiae* throughout a wine fermentation are consistent, which is reflected in a core gene expression program (CGEP) that is present across diverse fermentation conditions (1–4). However, this is not to say that *S. cerevisiae* metabolism is fixed, as *S. cerevisiae* dynamically responds to differences in the fermentation environment (e.g., nutrient levels, temperature, and differences in microbial communities) to maintain cellular metabolism and overall fitness (1, 2, 5, 6). For example, differences in grape must nitrogen concentrations leads to changes in metabolism that are accompanied by altered aroma compound in wine (7). This highlights the fact that metabolic adaptation to varied fermentation environments leads to differences in wine fermentation outcomes, including sensory differences. This relationship is mirrored by many findings showing that genetic changes causing altered expression of select genes or pathways in *S. cerevisiae* leads to quantifiable differences in wine fermentation outcomes (8). These facts support the generally accepted idea that interactions between *S. cerevisiae* and the unique chemical and biological matrix of each grape must are central to defining primary fermentation characteristics. We reason that these differences are the result of 1) the expression of unique genes outside those in the CGEP required for fermentation and/or 2) variation in expression of CGEP genes that changes the activity of various core pathways during fermentation.

The chemical and biological diversity of grape musts is in large part a reflection of biotic and abiotic pressures encountered by a grapevine during a growing season. For example, wines produced under similar vinification conditions from genetically identical grapes grown in different locations have diverse sensory outcomes (9), many of which are reproducible across multiple vintages (10). After observing diverse sensory outcomes in wines where a consistent variable between fermentations was vineyard location (9), we sought to uncover quantifiable contributions of vineyard site by using *S. cerevisiae* gene expression as a biosensor to detect differences between fermentations. This is motivated by the fact that high throughput gene expression surveys (microarray and RNA sequencing) have revealed the causes of stuck and sluggish fermentations (11), triggers for entry into stationary phase (1, 2), and the impact of inter-species interactions on *S. cerevisiae* metabolism in wine (6, 12, 13). In addition, as an organism commonly used in life science and biotechnology research, the *S. cerevisiae* genome and transcriptome is well understood with published datasets focused on gene expression in diverse environments, including wine (1–3, 14–16). This makes *S. cerevisiae* a powerful tool for understanding the wine fermentation environment and identifying key biotic and abiotic factors underlying fermentation outcomes.

Towards this end, we performed time series RNA sequencing on Pinot noir fermentations with the aim of identifying gene expression differences by vineyard site. However, using standard analysis methods (17–22), we only identified the dominant and consistent global shift in gene expression, the CGEP, across fermentations and not gene expression patterns indicating altered *S. cerevisiae* metabolism that would differentiate vineyard site (4). A major issue was that sampled fermentations proceeded at different rates, leading to asynchronous biological progression among sequenced samples with respect to fermentation status (e.g., sugar consumption). This was problematic because samples need to be at the same stage of fermentation to interpret the biological significance of differentially expressed genes (3, 23). This is a common problem in time series experiments with multiple groups and in some experimental systems there are strategies to combat this issue (23). For example, in experiments that study the cell cycle, inhibitors arrest the cell cycle at the same stage across groups thereby enabling comparisons (24). This experimental approach is not applicable to wine fermentations where nutrient availability is more important than stage of the cell cycle (25).

To address a similar issue, methods have recently been developed for the analysis of single cell RNA sequencing data from differentiating cells. In these experiments, as cells differentiate, absolute time may not reflect the extent of differentiation in each individual cell (26). Consequently, pseudotime analysis has been used to reorder cells from absolute time to the stage in differentiation relative to other cells undergoing the same process (26). In particular, diffusion maps have been used to reorder asynchronous cell populations because this analysis approach preserves relationships between samples (26). Diffusion mapping is a manifold learning technique that uses information from the *k* most similar samples to construct non-linear composites of the major sources of variation among samples (27, 28). Diffusion mapping has strengths over other dimensionality reduction algorithms, like principal component analysis (PCA) and t-distributed stochastic neighbor embedding (tSNE) that are applied to sequencing data to identify sources of variation (29, 30). The major advantages are that diffusion mapping is non-linear and robust to the “horseshoe effect” (unlike PCA) (31), preserves distances between samples (unlike tSNE), and is insensitive to sampling density. As a dimensionality reduction algorithm, diffusion maps extract latent variables that are inferred from relationships in the data, which can be used to represent composite sources of variation between samples.

Here, we use diffusion mapping to analyze time-series RNA-sequencing data from *S. cerevisiae* during wine fermentation. We use the resulting diffusion maps to synchronize gene expression across treatment groups and to extract latent variables, termed diffusion components (DCs), which represent the dominant sources of structure in the data. Diffusion maps *per se* provide no suggestion of the underlying genes that lead to separation of samples along diffusion components; therefore, we apply continuous differential expression analysis using each diffusion component to determine what genes vary among samples across a given diffusion component. Notably, this method was not exclusively useful for the analysis of RNA sequencing data from wine fermentations. We additionally apply this method to time series data collected from *S. cerevisiae* during hypoxia, which showed that diffusion mapping captures the dominant global shift in gene expression that occurs when yeast transitions from aerobic to anaerobic metabolism. These findings suggest that our method enables analysis of diverse asynchronous time series gene expression data, revealing both global shifts and subtle differences in gene expression among groups. In the context of wine, we find that diffusion mapping extracted the CGEP across Pinot noir fermentations, in addition to distinguishing subtler differences between fermentations that reflect differences in the grape musts. These findings offer important insights into variable wine fermentation and sensory outcomes driven by vineyard specific cell-environment interactions.

## Results and Discussion

Diffusion maps reorder asynchronous cell populations while preserving relationships between samples (26), which provides latent variables that represent non-linear composites of genes that vary between samples (**Figure 1**). We refer to these latent variables as *diffusion components*, the number of which is constrained by the number of samples in the data. Within each diffusion component (DC), a sample is represented by a single value and samples that have similar gene expression profiles among the genes captured in that component will have similar values. Moreover, samples at the origin of a DC (i.e., near 0) have gene expression profiles that do not vary along that component, while samples with positive or negative values diverge. Each DC captures diminishing structure among samples with the first diffusion component (DC1) accounting for the largest variation among all samples. Differential expression analysis can then be used to identify genes that vary among the samples along a given diffusion component. We expect the information extracted from each DC in this way will provide important insight into gene expression patterns caused by the time in fermentation (e.g., based here on Brix level) and vineyard site.

**Figure 1:**
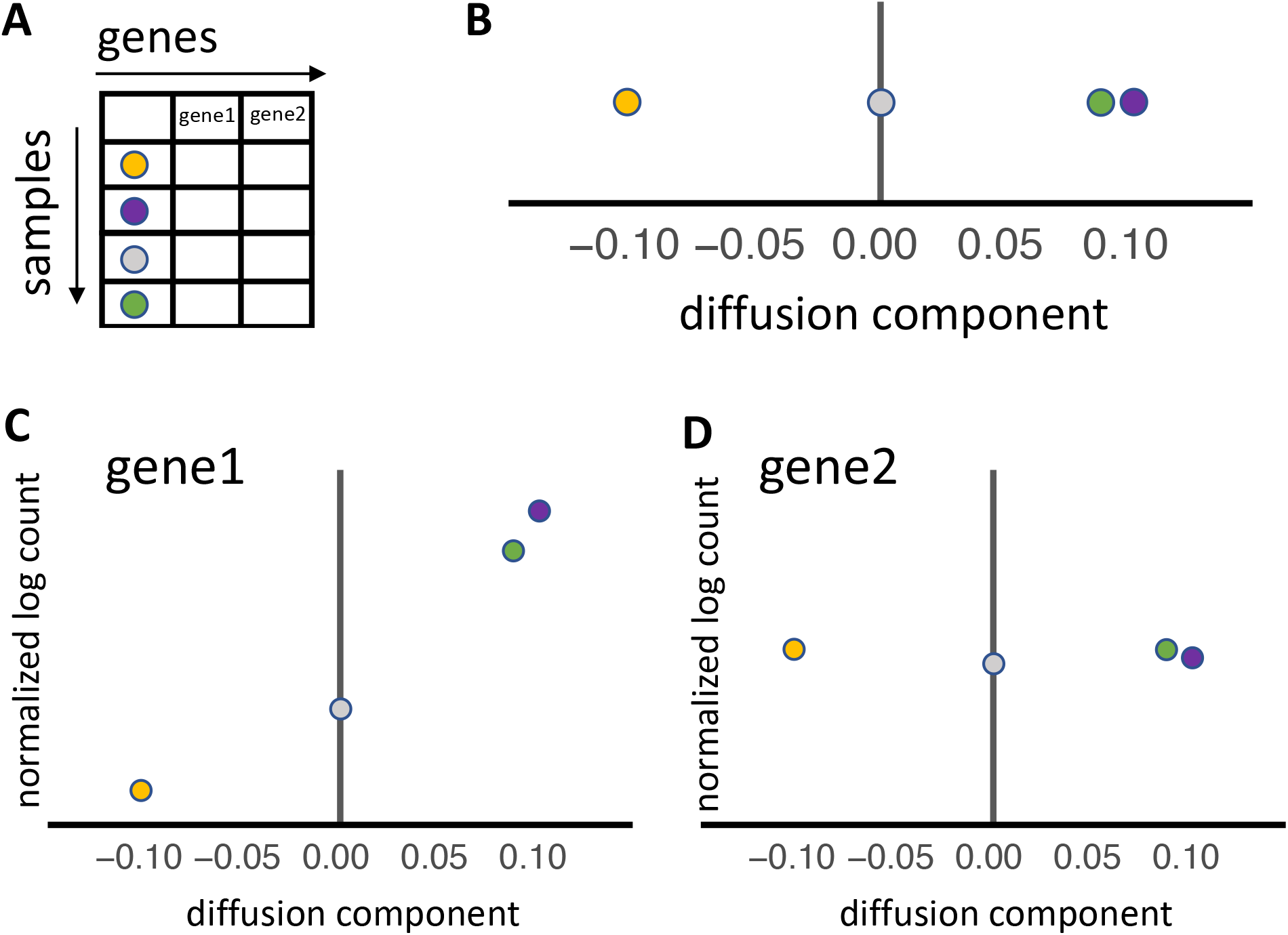
Extracting data from diffusion maps. Diffusion maps provide the underlying manifold in gene expression data through non-linear dimensionality reduction. **A**) When applied to many genes across many samples, diffusion maps extract features that represent combinations of genes that drive similarities and differences among samples. **B**) These features are termed diffusion components. Samples at either extreme of the diffusion component are the most different from each other, while samples that fall at the origin are invariant along that component. In the above graphic, the orange and purple dots are the most different, while the purple and green dot are most similar. Diffusion maps do not provide information on which genes lead to separation of samples along each diffusion component. **C**) Performing differential expression using the diffusion component as a continuous variable reveals the genes that significantly contribute to separation of samples. In the graphic, gene 1 has significantly higher expression in samples that fall on the right extrema of the diffusion component, as compared to samples that fall on the left extrema. **D**) While all genes are used to perform differential expression, not all genes are differentially expressed along an individual diffusion component. In this example, gene2 is not differentially expressed along the diffusion component.

### Diffusion mapping captures the global shift of gene expression during primary fermentation

We had previously performed inoculated primary fermentations of genetically similar Pinot noir grapes grown in 15 vineyard sites in California and Oregon over five vintages at the UC Davis Teaching and Research Winery (**Figure 2A**) (4, 9, 32, 33). We profiled the 2017 and 2019 vintages using time-course RNA sequencing data, with the aim of using *S. cerevisiae* gene expression as an indicator of similarities and differences across fermentations. In the 2017 vintage, we took samples approximately corresponding to early growth phase (16 hours), late growth phase (40 hours), stationary phase (64, 88 hours), and end of fermentation (112 hours) (**Figure 2B**). In the 2019 vintage, we shifted sampling to capture cellular adaptation after inoculation (2 and 6 hours) and reduced sampling later in fermentation (16, 64, and 112 hours). The initial grape musts varied in parameters like initial nitrogen, pH, malic acid, tartaric acid, non-*Saccharomyces* microbial profile, and elemental profile, while the final wines differed in volatile profiles and sensory characteristics (9, 32, 33). From these data we determined that *S. cerevisiae* had a consistent core gene expression program (CGEP) across fermentations from different vineyards and vintages (4).

**Figure 2:**
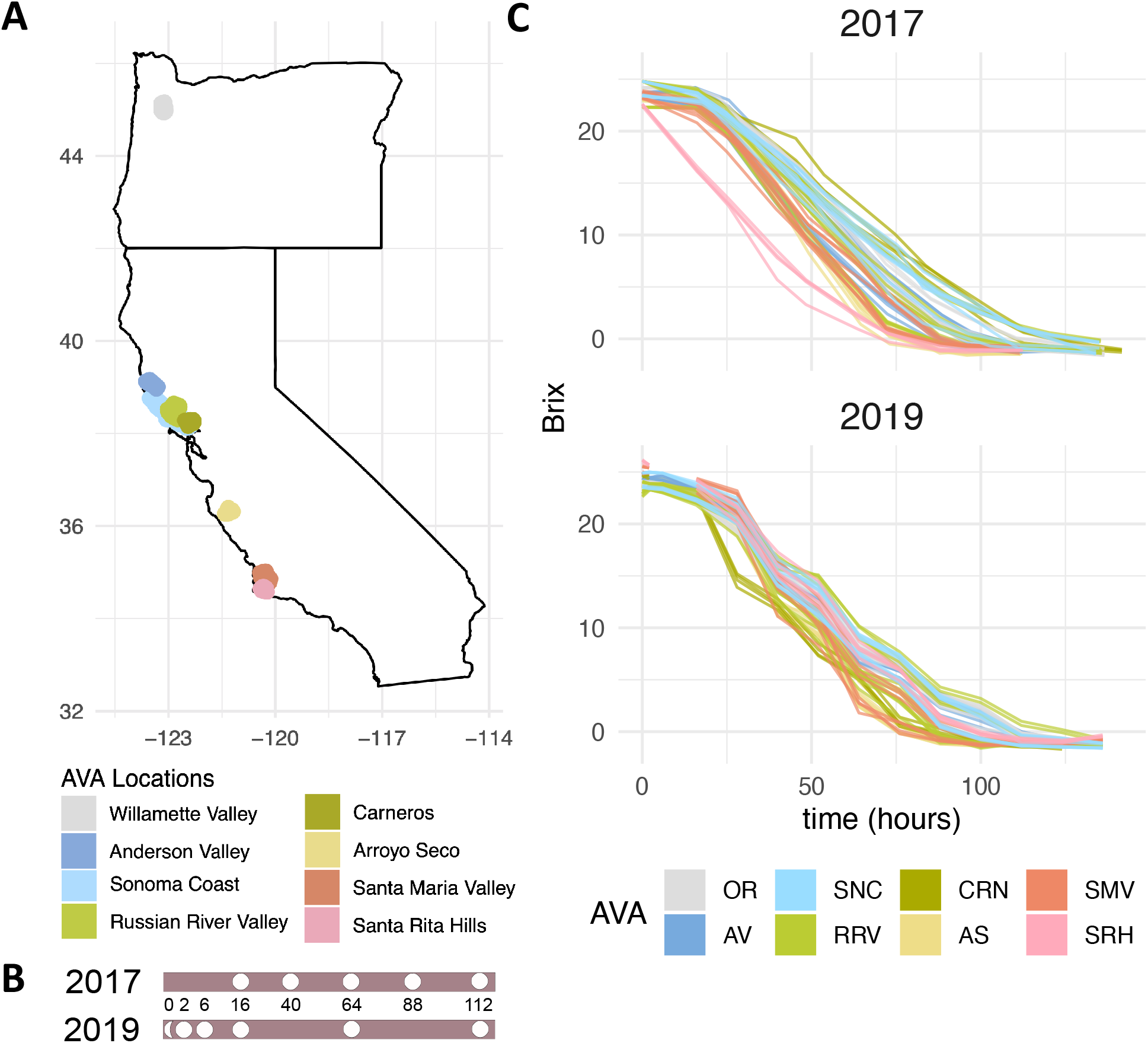
AVA locations of vineyard sites and sampling info across wine fermentations. Map of vineyard site American Viticultural Areas represented in this study. **B**) Primary fermentation sampling time points for 2017 and 2019 vintages. Times are shown in hours and are relative to inoculation. **C**) Vineyard site fermentation kinetics where Brix (total soluble solids) is used as a proxy for sugar levels.

Given the variable inputs and sensory differences described for wines from these vineyards (9), we expected that there would be differences in *S. cerevisiae* gene expression that may include genes known to impact the sensory outcome of wine (34). Yet, in our previous analysis (4), site-specific differences were hard to quantify because fermentations progressed at different rates, even with rigorous control of temperature at a 200 L scale, leading to asynchronous biological progression among samples with respect to sampling time (**Figure 2C**). To address this issue and gain insight into vineyard-specific factors altering fermentation outcomes, here we have applied diffusion mapping to extract DCs and performed differential gene expression analysis using DC values as a continuous variable to identify key gene expression patterns differentiating these fermentations.

In DC1, which accounts for the largest variation among all samples, we captured a clear transition during fermentation that is observed by the ordering of samples across DC1 in both the 2017 and 2019 vintages based on Brix (**Figure 3A-B**). In a previous analysis, we used Brix to perform continuous differential expression and identified the CGEP of *S. cerevisiae* in these Pinot noir fermentations (4). To test whether diffusion mapping and DC1 captured the CGEP during fermentation, we performed differential expression over DC1 and compared this to values calculated previously across the Brix variable (4). Log_2_ fold change values were strongly correlated between both methods of differential expression (**Figure 3C, Figure S1**), indicating that DC1 captured the dominant global shift in gene expression during both 2017 and 2019 fermentations.

**Figure 3:**
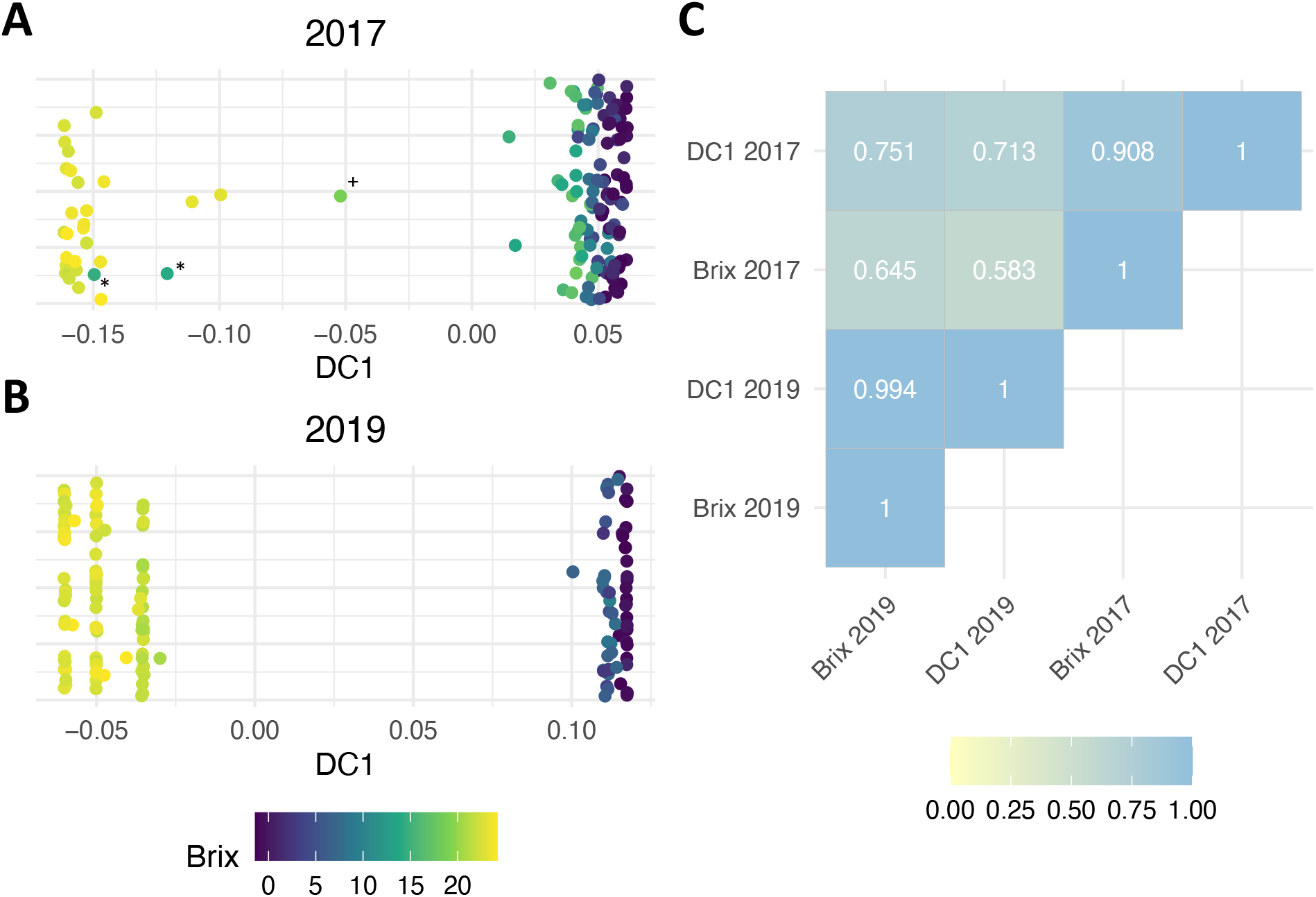
Diffusion mapping captures the core gene expression program of S. cerevisiae during wine fermentation. **A, B**) DC1 captures the metabolic transition that occurs as Brix decreases during fermentation. The same transition occurs in the 2017 vintage and the 2019 vintage. Each point represents a sample from one time from one fermentation. Points that are closer along the x axis are more similar. The y axis is jittered to allow all points to be visualized. Points are colored by Brix, a proxy for sugar concentration during fermentation, where Brix = 0 indicates end of fermentation. **C**) Graphic displays calculated correlations between differentially expressed genes in diffusion component 1 and genes that are differentially expressed as Brix decreases. Select samples in panel 3A are marked (* = SMV and + = OR sites) for ease of identification.

We next explored subsequent diffusion components (e.g., DC2 through DC8) to determine other less dominant drivers of structure among samples. In general, within the 2019 data set we observed that lower diffusion components (DC2-DC4) captured structure across samples relating to the time of fermentation (**Figure 4**). For example, we observed separations along DCs 2-4 based on Brix levels and not vineyard site (compare **Figure 4A and 4B**). These differences were driven by cellular remodeling in early fermentation (DC2, DC3) and starvation during late fermentation (DC4), based on the differentially expressed genes associated with each diffusion component (**Table S1**). In contrast, higher diffusion components (DC5–DC8) showed patterns that suggest these DCs varied more based on vineyard site than on time in fermentation (**Figure 5 and Table S1**). In addition, we saw the total number of differentially expressed genes diminished as the diffusion component number increased, indicating more subtle differences between samples (**Table 1**).

**Figure 4:**
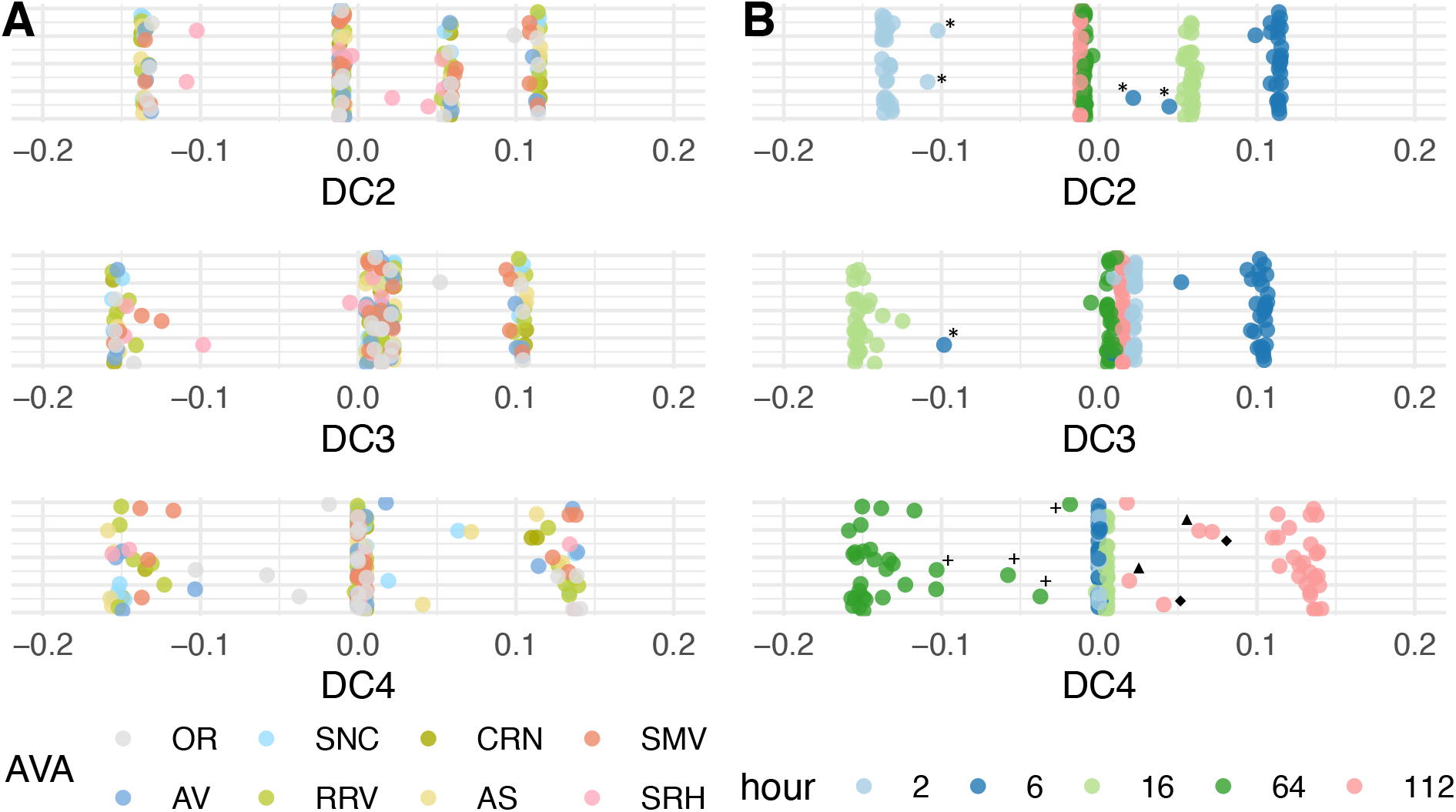
One-dimensional plots of DC2–DC4 for the 2019 vintage. Plots are colored by **A**) American Viticultural Area (AVA) and **B**) hours post inoculation and show that diffusion components capture different latent structure in the dataset. Each diffusion component represents a different relationship among samples with these components appearing to capture shifts between stages of fermentation, not vineyard site. Select samples in panel 4B are marked (* = SRH, + = OR, ▲ = SNC, and ◆ = AS sites) for ease of identification.

**Figure 5:**
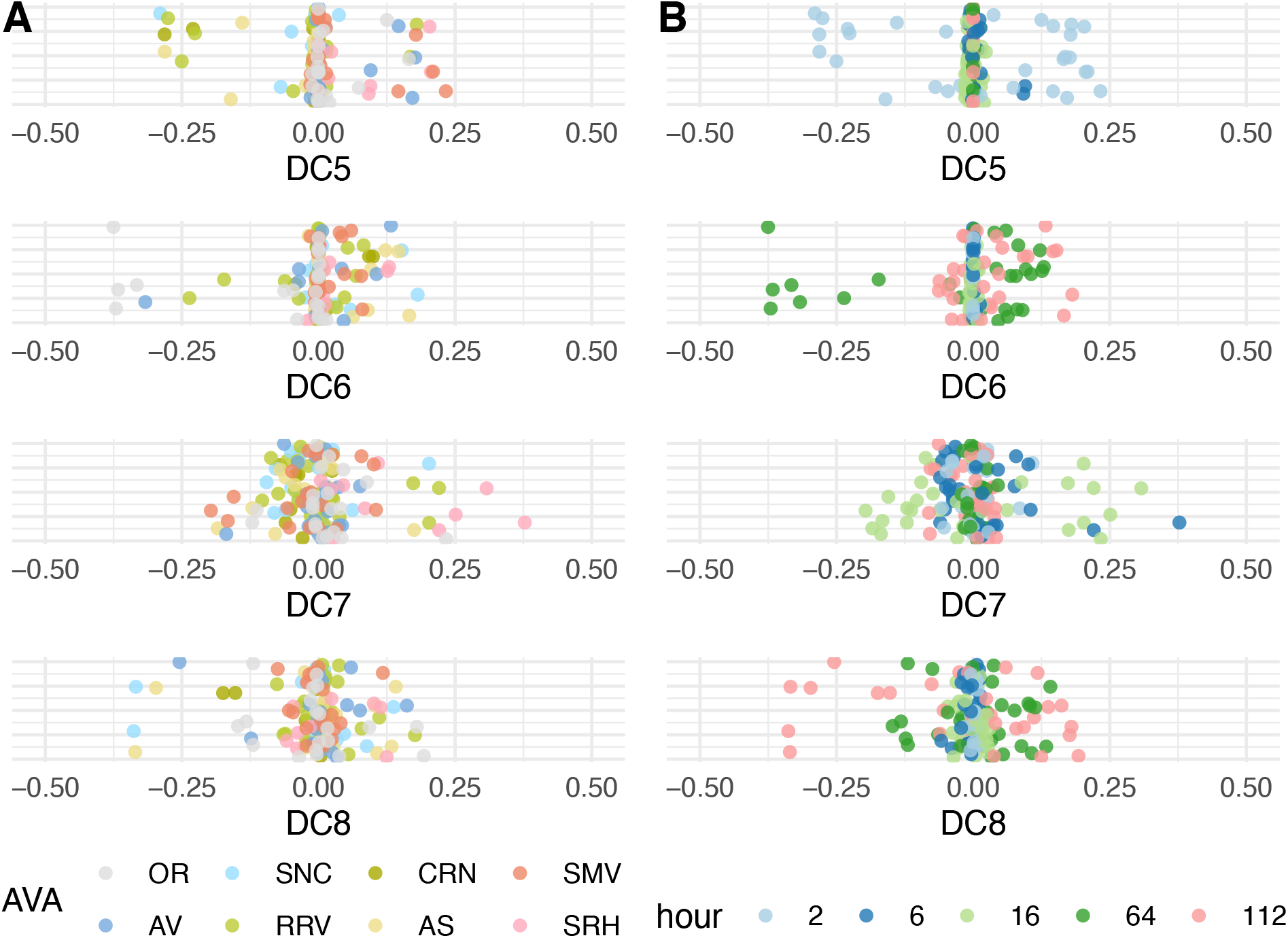
One-dimensional plots of DC5–DC8 for the 2019 vintage. Plots are colored by **A**) American Viticultural Area (AVA) and **B**) hours post inoculation. Higher diffusion components capture differences between vineyard site within a stage of fermentation, as seen by samples clustering together along a single or multiple diffusion components based on AVA, not Brix, indicating similarity in gene expression profiles within an AVA.

**Table 1:**
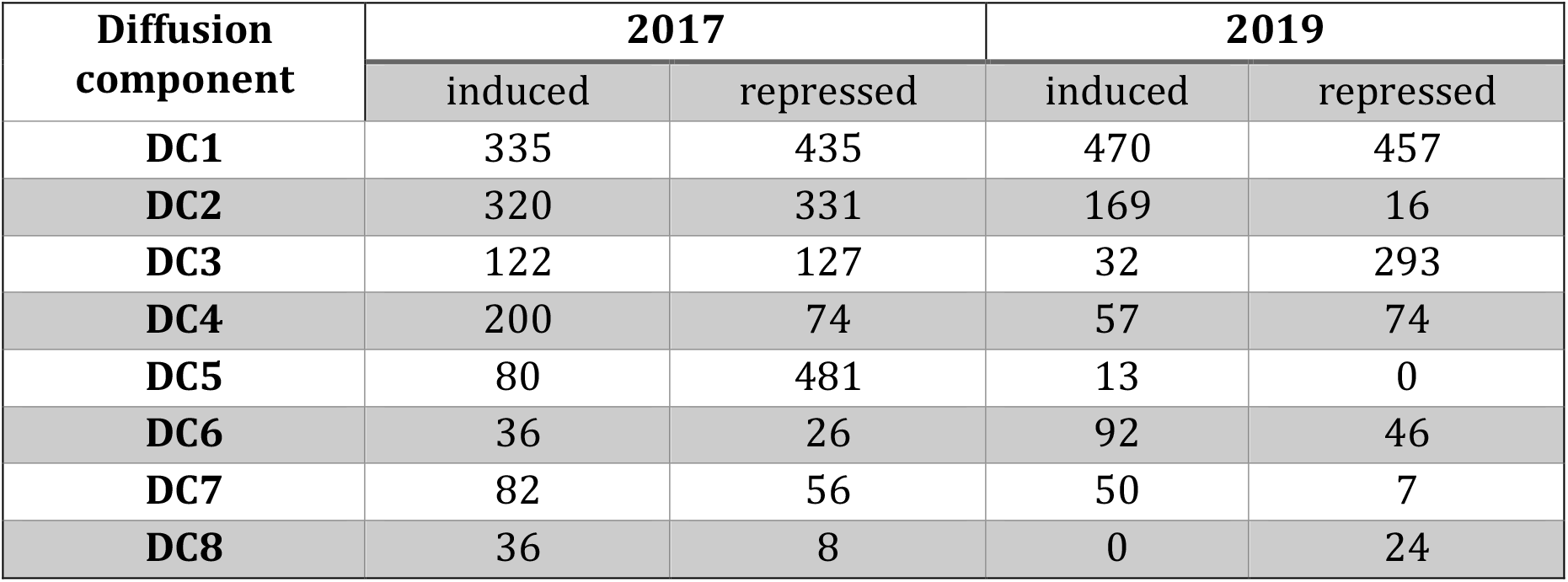
Number of significantly differentially expressed genes for the top diffusion components for the 2017 and 2019 vintages.

Notably, while we detected the global shift in gene expression during primary fermentation from data collected in 2017 along DC1 (**Figure 3A**), we did not observe clear delineations among higher DCs (**Figure S2**). All gene expression data are inherently noisy (35), which is continuously improved upon by technical and methodological advancements in sequencing. Applying these advances, we sequenced the 2019 vintage with unique molecular identifier (UMI) barcodes and were able to demultiplex sequencing lanes and remove PCR duplicates (36). With this improvement in data quality in the 2019 vintage over 2017, we expect that signals associated with higher diffusion components were better captured in the 2019 sequencing data. These improvements are also likely reflected in the number of differentially expressed genes across each DC in the 2017 vs. 2019 vintages (**Table 1**). Consequently, our are focused on the 2019 data.

### Diffusion mapping identified global shifts in gene expression during hypoxia

While diffusion maps successfully distinguished major fermentation transitions, we wanted to confirm that this method could consistently report on other RNA-sequencing data where global shifts in gene expression occur. To address this, we identified a publicly available gene expression dataset during hypoxia in *S. cerevisiae* (GSE85595 and GSE115171). Hypoxia occurs when a cell becomes oxygen limited, which is accompanied by large scale reprograming of gene expression for continued growth (37). We applied diffusion mapping to this dataset and observed an ordered time-dependent transition to a hypoxic phenotype along DC1 (**Figure 6A**). Sample positions along DC1 show a rapid transition within 5 minutes of nitrogen exposure, indicating a fast metabolic transition to hypoxia that matured over the remainder of the time course. As part of this genetic reprogramming, a transient shift in gene expression has been previously identified at ~30 minutes of the hypoxic response and shown to partially overlap with the environmental stress response (37). Within the diffusion mapping data, DC6 differentiated this transient state at 30 minutes of hypoxia (**Figure 6B**).

**Figure 6:**
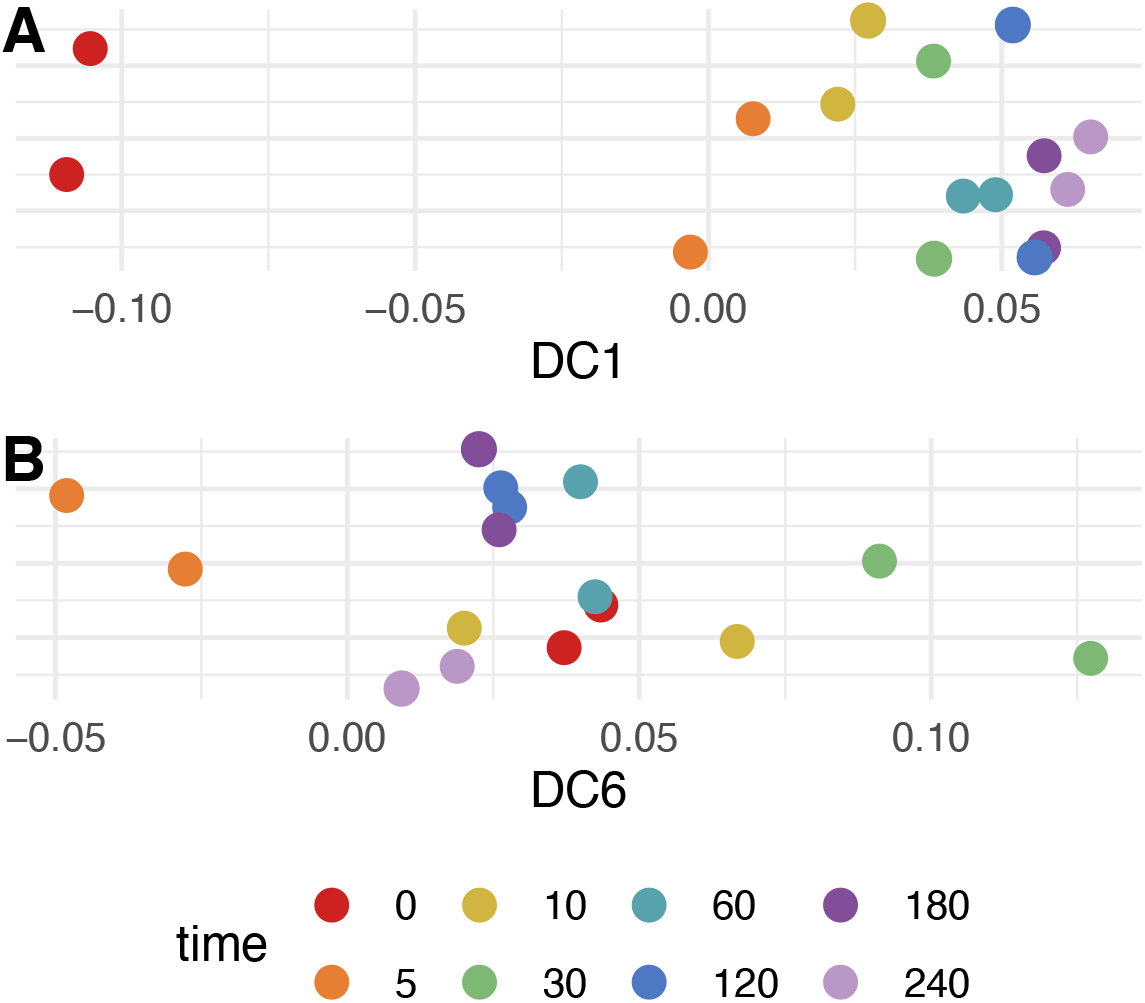
Diffusion map of S. cerevisiae exposed to nitrogen for 0–240 minutes. The trajectory of samples are displayed along **A**) DC1 that captured the transition from aerobic to anaerobic metabolism and **B**) DC6 that captured a transient transcriptome remodeling at 30 min.

We next investigated whether genes differentially expressed along DC1 matched oxygen-regulated genes identified by previous studies of hypoxia. Across seven microarray studies, 11 genes (3 aerobic, 8 hypoxic) were consistently identified as being involved in aerobiosis or anaerobiosis in at least six studies (compiled by Bendjilali et al. in (37)). We identified all 11 of these genes as differentially expressed (p < 0.05) along DC1. We further compared our results to time-series RNA-sequencing profiles of wild type *S. cerevisiae* during a hypoxic response (37). Along DC1, 239 of 291 (82.1%) aerobic genes were significantly expressed prior to exposure to nitrogen, while 422 of 519 (81.3%) hypoxic genes were significantly induced after prolonged exposure to nitrogen (**Table S2**). Genes expressed at time zero were enriched in pathways like ribosome biogenesis, oxidative phosphorylation, and sterol metabolic process, while genes induced after prolonged exposure to nitrogen were enriched in pathways like oxidation reduction process, cell wall, glycogen metabolic process, and glycolysis/gluconeogenesis (**Figure S3**). These findings align well with our knowledge of the hypoxic transition in yeast (37, 38). Moreover, these results indicate that diffusion mapping accurately captured global changes in gene expression during the hypoxic shift, including transient gene expression states.

### Diffusion mapping detected metabolic remodeling throughout wine fermentation

With the knowledge that diffusion mapping effectively identified gene expression changes across two different data sets, we set out to investigate the gene expression patterns captured within each DC in our wine fermentation datasets. In the 2019 data, along DC2 there were clear separations among the 2, 6, and 16 hour samples, while the 64 and 112 hour samples fell on the origin (**Figure 4B**). Within the genes captured along this component, arginine biosynthetic process was enriched in genes that were more highly expressed in the 2 hour samples *(ARG1, ARG3, ARG5,6, ARG8)* (**Figure S4, Figure S5**). Arginine is likely the most abundant amino acid in Pinot noir grape must (39) and genes that encode proteins involved in arginine biosynthesis are suppressed by the presence of arginine (40). Expression of these biosynthetic genes in early fermentation likely reflects that *S. cerevisiae* is yet adapted to the wine environment by 2 hours after inoculation. By 6 hours of fermentation, expression of these genes decreases, potentially signaling completion of cellular adaptation to the grape must environment. Interestingly, four of the 16 genes (YMR244W, YPR078C, YGL117W, YER085C) expressed in the 2 hour samples have no known function. Given that very few genes were differentially expressed at 2 hours and they were enriched for arginine biosynthesis, one speculation is that these genes may have functions related to nitrogen and arginine biosynthetic processes. Alternatively, expression of these genes may be associated with other cellular processes associated with early adaptation to the must environment.

The 6 hours samples segregated to the opposite extrema of DC2 and were the most differentiated from the 2 hours samples along this component (**Figure 4B**). Glycolysis was enriched among genes induced in these samples (**Figure S5**), which was also accompanied by gene expression changes that support a transition to anaerobic metabolism. For example, we detected induction of anaerobic translation elongation factor encoded by *ANB1* (**Figure S4**), which is optimally expressed below 0.5 μmol/L O_2_ (41), likely indicating must oxygen levels at this timepoint. Genes important for cell wall processes were also induced at 6 hours, with *TIR1-4* being four of the top five genes induced (**Figure S4**). These genes encode cell wall mannoproteins that are required for anaerobic growth (42). These genes are also important in DC3 for separating the 6 and 16 hours samples, along with many genes induced by anaerobiosis that included *DAN1* and *PAU* genes (*PAU2-PAU5, PAU7, PAU8, PAU10-PAU12, PAU15-PAU17, PAU19, PAU20, PAU23*, and *PAU24*) in the 16 hour samples (43, 44). Together, the induction of these genes regulated in response to oxygen across DC2 and DC3 likely signal the transition to anaerobiosis. In DC3, there were also many other biological processes, cellular compartments, and molecular functions enriched among the 293 genes that were induced in the 16 hour samples (**Figure S6**), consistent with a transition to an active growth phase at this stage of fermentation. As diffusion components are ordered with the most variation among samples occurring first, DC2 and DC3 demonstrated that early metabolic remodeling was second only to gene expression changes that occur as Brix decrease (e.g., captured in DC1) during fermentation.

Along DC4, we observed separation of the 64 hour samples from the 112 hour samples. In the 64 hour samples, transmembrane transport, including amino acid and polyamine transport, were enriched categories among the genes that were induced (**Figure S7**). Many induced genes *(DUR3, DAL5, DAL7)* are involved in allantoin metabolism, a non-preferred nitrogen source (**Figure S4**). Induction of these genes at 64 hours likely indicates relief of nitrogen catabolite repression consistent with decreasing nitrogen concentrations and nutrient availability. Genes repressed by the presence of amino acids were also induced in the 112 hours samples *(GAT2, ARG3)*. Interestingly, *HXT13* and *MAN2* were among the top induced genes, along with *HXT17*, in the 112 hours samples (**Figure S4**). These two *HXT* genes encode mannitol transporters (45) and *MAN2* encodes mannitol dehydrogenase. Expression of these genes would enable *S. cerevisiae* to metabolize mannitol as a non-preferred carbon source (45–47). Mannitol is produced by *non-Saccharomyces* organisms, including lactic acid bacteria (48) and other non-*Saccharomyces* yeast (49). Expression of these genes late in fermentation likely signals an increasingly generalized metabolic program that is utilizing non-preferred carbon sources as preferred sugars were exhausted.

Overall, we interpret the patterns of separation along DC1-4 to be reflective of gene expression changes occurring as *S. cerevisiae* proceeds through fermentation, adapts to the increasingly nutrient limited environment, and deals with associated stresses. While these changes appear common to the fermentations conducted here, future work will be required to address if individual processes captured in DC2-4 occur in the context of other wine strains and grape varieties or are unique to the wine yeast RC212 and Pinot noir fermentations. Nonetheless, these observations indicate that this analysis approach is a robust means for dealing with asynchronous gene expression data across fermentations. Moreover, it raises many questions about the genes important for defining separation along these DCs, including gene products involved in arginine, mannitol, and anaerobic metabolism. Of notable interest are the large family of *PAU* genes, the vast majority of which have no known function in *S. cerevisiae*, but have been previously noted for being induced during fermentation and in response to stress (50).

### Lower diffusion components captured varied progression through fermentation

Along lower diffusion components, patterns of separation were based on the time of fermentation (discussed above), but outliers from select vineyard sites were also noted. We expect this indicates site-specific differences that influence *S. cerevisiae* activities during fermentation. For example, in the 2017 vintage, samples taken at 40 hours from SMV sites and OR1 were shifted toward the 16 hour samples along DC1 indicating a different metabolic trajectory (**Figure 3**). Interestingly, we detected *Lactobacillus kunkeei* transcripts in primary fermentations from SMV sites (4, 32). *L. kunkeei* is a fructophilic lactic acid bacteria that produces mannitol and lactic and acetic acids during wine fermentation (48). The observation of an altered metabolic program in early fermentation with the unique presence of *L. kunkeei* may indicate inter-species interactions that modulated *S. cerevisiae* gene expression. In support of this, expression of genes involved in mannitol metabolism at 112 hours were the highest for the SMV sites *(HXT13, HXT17, MAN2)*. One possibility is that *L. kunkeei* may have induced a [GAR+] prion state, which is known to alter the metabolic strategy employed by *S. cerevisiae* (51). Following these observations in 2017, we attempted to detect the [GAR+] prion state in yeast isolated from the 2019 fermentations, but failed to detect [GAR+] yeast.

Other outliers included SRH1, which in the 2019 vintage at 6 hours, and to a lesser extent at 2 hours, did not cluster with other samples from these same time points along DC2 and DC3 (**Figure 4**). The SRH1 6 hour samples were instead shifted toward other 16 hour samples, potentially indicating faster cellular adaptation to the wine environment. In support of this, fermentations from SRH sites decreased in Brix faster than other sites in the 2017 vintage (**Figure 2C**). Another example involved 64 hour samples from OR1 and OR2 sites being shifted along DC4 toward the 112 hour cluster (**Figure 4**), which may relate to nutrient conditions specific to OR sites (see further discussion below). Similarly, 112 hour samples from SNC1 and AS2 were shifted toward the 64 hour cluster (**Figure 4**). We expect these outliers in the lower DC components related to the time in fermentation reflect differences between the musts (e.g., nutrient levels or presence of specific non-*Saccharomyces* organisms) that impact *S. cerevisiae* metabolism and the timing of gene expression transitions as fermentations progress.

### Higher diffusion components identified vineyard specific gene expression patterns

The common patterns and existence of outliers across lower diffusion components, which aligned with fermentation progression and presence of non-*Saccharomyces* species, indicated that information about specific sites were captured by these analyses. Given that some higher diffusion components clearly separated samples within a stage of fermentation **(Figure 5)**, we used gene expression differences across the higher DCs to investigate vineyard-specific patterns (**Figure S8** and **Table S1**). In this way, we aimed to identify *S. cerevisiae* activities, inferred by the genes involved, specific to a vineyard site(s). We specifically concentrated on the 2019 samples that separated to the extremes of each DC, as this separation indicates that these samples were the most differentiated at the transcriptome level. For example, at 2 hours SMV1, SMV2, SRH1, AV2, and RRV3 were most separate from RRV2, CRN1, SNC1, and AS2 along DC5 (**Figure 5**). When comparing these sites, a standout difference was the induction of genes in samples from SMV1, SMV2, SRH1, AV2, and RRV3 that were enriched for vitamin metabolic and cell wall processes (**Figure S8A and S9**). Given that co-culture experiments have demonstrated that *S. cerevisiae* induces genes involved in cell wall remodeling and vitamin biosynthesis in response to the presence of *non-Saccharomyces* yeasts (6, 52, 53), we tested whether detected gene expression by *non-Saccharomyces* yeasts in 2 hour samples correlated with DC5. Indeed, DC5 values correlated with total gene expression of *Hanseniaspora uvarum* (R^2^ = 0.49, p < 0.001), but not with total gene expression of other organisms (**Table 2**), suggesting that *H. uvarum* activity leading up to these early fermentation samples may have impacted *S. cerevisiae* metabolism. This is consistent with a previous study which reported that *S. cerevisiae* remodels its cell wall in the presence of *H. uvarum* at three hours post-inoculation in a wine fermentation (6). Interestingly, *PDC5* was among genes induced in fermentations with higher gene expression by *H. uvarum* along DC5 (**Table S1**). *PDC5* encodes one of three isoforms of pyruvate decarboxylase, an enzyme involved in the formation of flavor-active higher alcohols in wine via the Ehrlich pathway (54). This suggests that the presence of *H. uvarum* may lead to gene expression changes that directly impact wine sensory outcomes. The activity of this non-*Saccharomyces* yeast may also relate to the altered metabolic program detected in SMV1/2 and SRH1 fermentations within the lower diffusion components (as discussed above). Notably, in a previous study we found that all fermentations had detectable *H. uvarum* DNA, but only a subset had detectable *H. uvarum* gene expression (32). Given the potential for *H. uvarum* to impact *S. cerevisiae* gene expression and metabolism, in the future it will be important to determine the factors that lead to *H. uvarum* (in)activity in select fermentations.

**Table 2:** Correlation between total gene expression by non-Saccharomyces organisms and DC5.

| Organism | R<sup>2</sup> | p value |
| --- | --- | --- |
| *Aureobasidium pullulans* | -0.03555 | 0.947 |
| *Botrytis cinerea* | -0.03261 | 0.774 |
| *Cladosporium sp SL 16* | -0.03341 | 0.804 |
| *Hanseniaspora opuntiae* | -0.03524 | 0.911 |
| *Hanseniaspora uvarum* | 0.490605 | < 0.001 |
| *Lachancea thermotolerans* | -0.02741 | 0.638 |
| *Metschnikowia fructicola* | 0.069637 | 0.086 |
| *Pichia kudriavzevii* | -0.02819 | 0.654 |
| *Rhizopus stolonifer* | -0.02439 | 0.582 |

SMV and SRH are neighboring American Viticultural Areas in southern California (**Figure 2A**). While samples from the SMV sites and SRH1 group together at 2 hours, we detected separation of these fermentations at 16 hours along DC7 (**Figure 5**). This suggests that while these sites were initially similar, they differed later in fermentation. While few genes were significantly induced in SMV vs. SRH samples along DC7, *ADH4* was the top induced gene (**Figure S8C**). *ADH* genes encode alcohol dehydrogenases that play an important role in fermentation by facilitating transitions between acetaldehyde and ethanol involving the redox cofactor NAD+. *ADH1* encodes the primary alcohol dehydrogenase isoform responsible for this reaction during wine fermentation (55). Alcohol dehydrogenases are also involved in the formation of fusel alcohols within the Ehrlich pathway (56). As such, differences in *ADH4* gene expression could be an important site-specific difference that has a role in *S. cerevisiae* metabolism and wine aroma development. Other genes more highly expressed in SMV sites were involved in cell growth processes, including translation *(MRP2* and *TIF2)*, transcription *(MED1*), and cell division *(CLB6)* (**Figure S8C**). In site SRH1, more highly expressed genes along DC7 vs. SMV were involved in oxidative stress *(RCK1)* and sporulation *(SPO74* and *SSP1)*. These site-specific differences in gene expression involving growth vs. stress indicate varied fermentation environments leading to altered gene expression at 16 hours. Given that genes associated with the Ehrlich pathway and fusel alcohol anabolism differentiated SMV sites and SRH1 at 2 and 16 hours of fermentation, the Ehrlich pathway may be an important component to consider in the context of site-specific differences in these Pinot noir wines.

In the case of DC6, at 64 hours OR1/2 and RRV2 samples segregated to one extrema (**Figure 5**). Genes induced in these samples were associated with nitrogen limitation *(DAL5, PUT1, PUT2;* (1)), while genes involved in ammonia metabolism *(MEP3, SSY1, AUA1)* were induced within fermentations from sites at the other extrema (**Table S1**). In line with these patterns that reflect differences in nitrogen availability, DC6 values correlated with initial grape must nitrogen as measured by an o-phthaldialdehyde assay (NOPA) and NH3 measurements (initial NOPA: R^2^ = 0.62, p < 0.001, initial NH3: R^2^ = 0.60, p < 0.001), led by low initial nitrogen levels in OR1, OR2, and RRV2 (**Figure 7**). Interestingly, while initial nitrogen levels in OR1, OR2, and RRV2 were the lowest among all fermentations, these fermentations were supplemented approximately 24 hours after inoculation with a combination of DAP and a complex nitrogen sources to adjust total yeast assimilable nitrogen (YAN) levels to 250 mg/L. Yet, these data indicate persistent nitrogen limitation for these sites, suggesting that these nitrogen additions may not have been sufficient to meet nutrient requirements in these fermentations. These findings suggest that more research is needed to understand the impact of nitrogen additions on fermentation, including the timing of addition and the nitrogen source.

**Figure 7:**
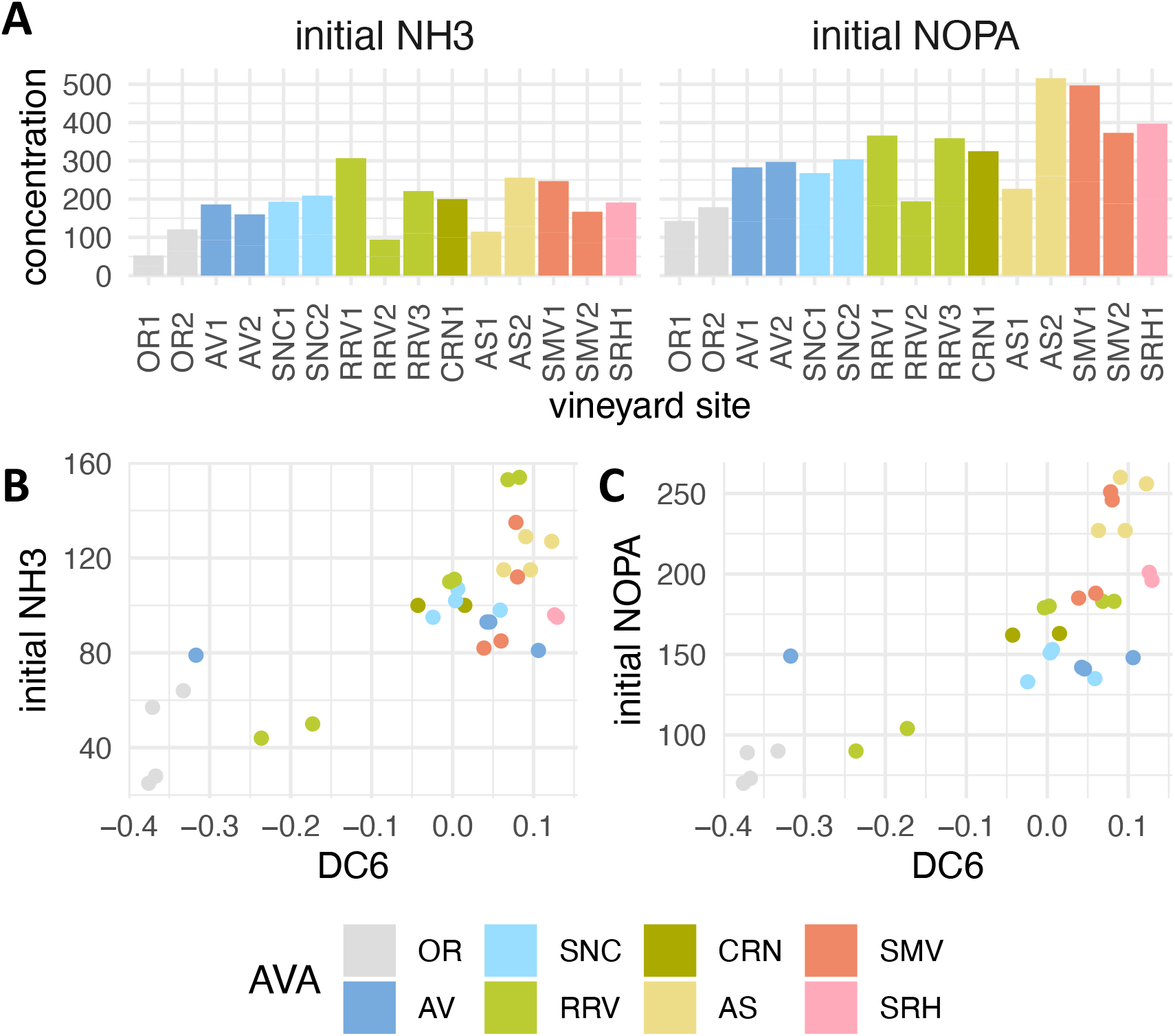
DC6 separates samples by initial nitrogen concentration in the grape must. **A**) Initial concentration of NH_3_ and nitrogen by o-phthaldialdehyde assay (NOPA) was lower in fermentations from OR1, OR2, and RRV2 than other vineyards. **B, C**) Initial NH3 and NOPA in grape must correlates with DC6, driven by low nitrogen in fermentations from vineyard sites OR1, OR2, RRV3.

DC8 separated 112 hour samples with SNC1 and AS2 segregating from other samples (**Figure 5**). Interestingly, 14 of the 24 genes induced in these samples are of unknown function (**Table S1 and Figure S8D**). Among the induced genes with known functions were *DDR2* and *HSP30*, which are stress related genes transcribed in response to a variety of environmental or physiological stresses (57), as well as *YDL218W* that is induced in response to the mycotoxin patulin produced by a variety of molds (58). Associated with these stress-related genes were genes that function in meiosis and sexual reproduction, including *SPO74, MFA1*, and *AFB1*. These data are suggestive of stresses in these fermentations that could be driving the diploid wine yeast into meiosis and a sexual reproduction cycle. This is of particular note, since the stresses associated with a wine fermentation environment are thought to impart strong selective pressures that drive adaptive evolution (59). This is reflected by the fact that *S. cerevisiae* strains associated with wine show a propensity for genetic diversity, including many instances of hybridization (60). Future research will be required to understand what particular stresses in SNC1 and AS2 are driving these unique patterns of gene expression, in addition to what outcome this has on fermentation performance.

Finally, across diffusion components, it is worth noting that fermentations from the same American Viticultural Area (AVA) were commonly grouped together (**Figure 4, Figure 5**). For example, fermentations from OR, AV, and SMV sites often grouped together, providing support for the concept of AVA and regional differences from the perspective of *S. cerevisiae* gene expression. However, we did not observe grouping among all fermentations from the same AVA along all diffusion components. For example, samples from AS grouped together along DC5 (2 hours) and DC6 (64 hours), but not in DC8 (112 hours). The AS sites are separated by 1 km and yet separation along DC8 suggests there was detectable variation in *S. cerevisiae* metabolism in primary fermentation (**Figure 4, Figure 5**). Replicates from these same vineyard site co-cluster, suggesting that lack of reproducibility in fermentations was not a factor in this observation. Similarly, fermentations from RRV did not cluster together along any diffusion component, suggesting that sub-appellations within the Russian River Valley are associated with significantly different *S. cerevisiae* gene expression patterns (**Figure 4, Figure 5**). This matches recent findings that show sub-regional variation in elemental profiles of wine from the Russian River Valley (61). Importantly, the gene expression differences we detected across each of these diffusion components provides candidate genes and pathways that may underlie vineyard-specific fermentation outcomes.

## Conclusion

In this study, we paired diffusion mapping with differential expression and captured global shifts in gene expression. This method revealed differences in primary fermentation of Pinot noir wine from 15 vineyard sites, as well as changes in *S. cerevisiae* gene expression induced by hypoxia. Diffusion mapping was especially well suited for these studies as in both cases cells progressed asynchronously through transcriptome changes with respect to sampling time. Through our analysis of wine fermentations, we found vineyard-site specific gene expression patterns that correlated with *H. uvarum* metabolic activity and initial nitrogen composition of grape must, as well as indications of sexual reproduction in select fermentations. Together, these data provide important insight into the wine fermentation environment, including pathways, genes, and environmental factors that should be considered in the context of differential fermentation outcomes. Given the tremendous complexity of gene-environment interactions, we expect these data also serve to highlight the large amount of work to be done to understand both the biological mechanisms at play and how this knowledge can be applied by industry. In particular, we report transcriptomic heterogeneity that arises from the same strain of yeast, fermented in the same facility, using grape must from genetically identical plants. How this variability changes across the diverse landscape of wine yeasts strains and fermentation environments (e.g., grape varieties and associated chemical and microbiological profiles) remains to be seen. Importantly, we expect that the approaches pioneered here for studying *S. cerevisiae* gene expression in a complex environment using diffusion mapping provide an effective tool to probe these questions.

## Methods

### Sampling, sequencing and preprocessing of wine fermentation samples

The winemaking protocol has been described previously (9, 62). The grapes used in this study originated from 15 vineyards in eight American Viticultural Areas in California and Oregon, U.S.A. All grapes were Vitis vinifera L. cv. Pinot noir clone 667, with rootstock 101-14 (AV1, RRV1, SNC1, SNC2, CRN1, AS1, AS2, SMV1, SMV2, SRH1), Riparia Gloire (OR1, OR2), or 3309C (AV2, RRV2, RRV3). Grapes were harvested at approximately 24 Brix and transported to University of California, Davis Pilot Winery for fermentation. We performed separate fermentations for grapes from each site, with two fermentations per site, totaling of 20 fermentations per vintage (40 fermentations total). After harvest, the fruit was separated into half-ton macrobins on harvest day and Inodose SO2 added to 40 ppm. Bins were stored in a 14°C cold room until destemming and dividing of the fruit into temperature jacket-controlled tanks. N2 sparging of the tank headspace was performed prior to fermentation and tanks sealed with a rubber gasket. We cold soaked the must at 7°C for three days and adjusted TSO_2_ to 40 ppm on the second day. After three days, the must temperature was increased to 21°C and programmed pump overs were used to hold the tank at a constant temperature. We reconstituted *S. cerevisiae* RC212 with Superstart Rouge at 20 g/hL and inoculated the must with 25 g/hL of yeast. At approximately 24 hours after inoculation, nitrogen content in the fermentations was adjusted using DAP (target YAN – 35 mg/L – initial YAN)/2, and Nutristart using 25 g/hL. Nitrogen was adjusted only if YAN was below 250 mg/L. Approximately 48 hours after fermentation, fermentation temperatures were permitted to increase to 27°C, and again added DAP using the formula (target YAN - 35 mg/L - initial YAN)/2, and fermentation were then continued until Brix < 0. Fermentation samples were taken for Brix measurements and RNA isolation at 16, 64, and 112 hours relative to inoculation. To ensure uniform sampling, a pumpover was performed ten minutes prior to sampling each tank. For RNA samples, 12mL of juice was obtained, centrifuged at 4000 RPM for 5 minutes, supernatant was discarded, and the cell pellet snap frozen in liquid nitrogen. Samples were stored at −80°C until RNA extraction.

### RNA Extraction and Sequencing

Yeast pellets were thawed on ice, resuspended in 5ml Nanopure water, centrifuged at 2000g for 5min, and aspirated the supernatant. RNA was extracted using the Quick RNA Fungal/Bacterial Miniprep kit including DNase I column treatment (cat#R2014, Zymo Research). RNA was eluted in 30μL of molecular grade water and assessed for concentration and quality via Nanodrop and RNA gel electrophoresis. Sample concentrations were adjusted to 200ng/μl and used for sequencing. 3′ Tag-seq single-end sequencing (Lexogen QuantSeq) was applied in both the 2017 and 2019 vintage, with the addition of UMI barcodes in 2019. The University of California, Davis DNA Technologies Core performed all library preparation and sequencing.

Sequencing samples were preprocessed according to manufacturer recommendations. First, we hard-trimmed the first 12 base pairs from each read and removed Illumina TruSeq adapters and poly A tails. Next, STAR was used to align reads against *S. cerevisiae* S288C genome (R64, GCF_000146045.2) with parameters --outFilterType BySJout --outFilterMultimapNmax 20 --alignSJoverhangMin 8 --alignSJDBoverhangMin 1 --outFilterMismatchNmax 999 --outFilterMismatchNoverLmax 0.6 --alignIntronMin 20 --alignIntronMax 1000000 --alignMatesGapMax 1000000 --outSAMattributes NH HI NM MD --outSAMtype BAM SortedByCoordinate (63). For the 2019 vintage, UMI tools was used to deduplicate alignments (64). Reads mapping to each open reading frame were quantified using htseq count (65).

### Hypoxia data set

Gene expression counts data was downloaded using from GEO using accession numbers GSE85595 and GSE115171.

### Construction of Diffusion Maps

Diffusion maps were built as described previously (66). To build diffusion maps from wine fermentations samples, k = 10 nearest samples was used, while for hypoxia k = 20 was used. We increased the k size for hypoxia given the larger number of samples (n = 146 in 2017 vintage, n = 150 in 2019 vintage, and n = 336 in hypoxia). Prior to diffusion map construction, gene counts were to non-mitochondrial mRNA and read counts normalized based on total number of reads per sample (library size).

### Differential expression

To determine which genes drove separation of samples along each component, differential expression was used to correlate each gene with diffusion component values. The R package limma was used to fit a linear regression model to each gene (20). As input to differential expression, raw sequencing counts were used as input to differential expression, and were filtered and normalized with the limma package using the calcNormFactors() function (20). Using this model, the log_2_ fold change is the slope of the line for each unit increase in the diffusion component. Log_2_ fold change values were normalized by calculating the length of the diffusion component and multiplying all log_2_ fold change values by this amount, (max - minimum) * log_2_ fold change. Log_2_ fold change values that were greater than two were analyzed, i.e., genes with a log_2_ fold change of at least 2 between the most separated samples along a diffusion component. Gene Ontology and KEGG enrichment analysis were performed using the R clusterProfiler package (67).

### Data Availability

Data is available in the Sequence Read Archive under accession PRJNA680606. All analysis code is available at github.com/montpetitlab/Reiter_et_al_2020_DiffusionMapping.

## Acknowledgements

We thank all past and current members of the Montpetit, Runnebaum, and Lab for Data Intensive Biology (Brown) laboratories for their support of this work. T.R. and C.T.B were supported by the Gordon and Betty Moore Foundation’s Data-Driven Discovery Initiative [GBMF4551]. T.R. was supported by the Harry Baccigaluppi Fellowship; Horace O Lanza Scholarship; Louis R Gomberg Fellowship; Margrit Mondavi Fellowship; Haskell F Norman Wine & Food Fellowship; Chaîne des Rôtisseurs Scholarship; Carpenter Memorial Fellowship. The authors would like to recognize support from Jackson Family Wines, in addition to support from Lallemand Inc.

---

**Figure S1:**
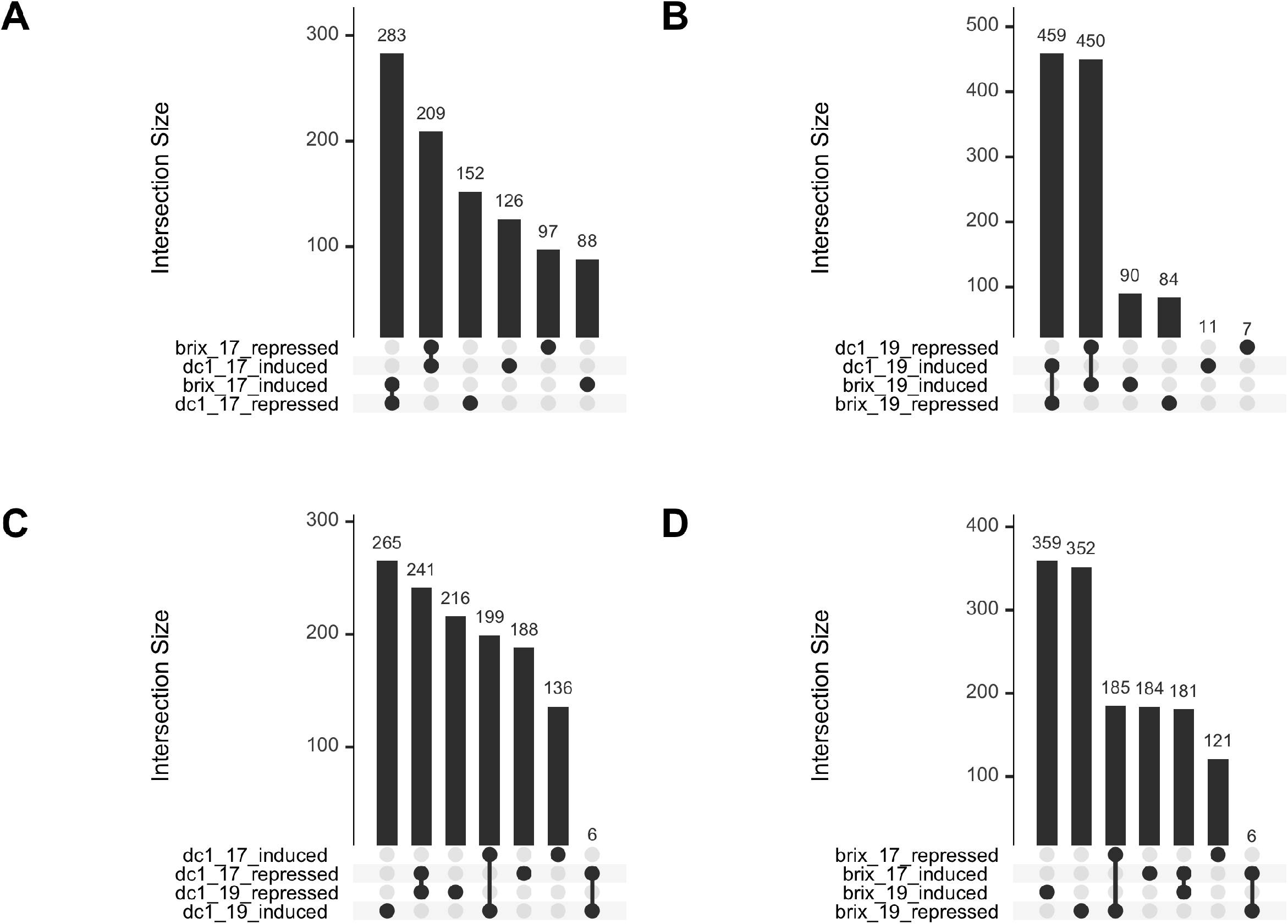
Upset plot depicting intersections between differentially expressed genes using diffusion component 1 or Brix as continuous variable. A) Intersection of differentially expressed genes from the 2017 vintage. **B**) Intersection of genes from the 2019 vintage. **C**) Intersection of genes differentially expressed along diffusion component 1 from the 2017 and 2019 vintages. **D**) Intersection of genes differentially expressed relative to change in Brix from the 2017 and 2019 vintages.

**Figure S2:**
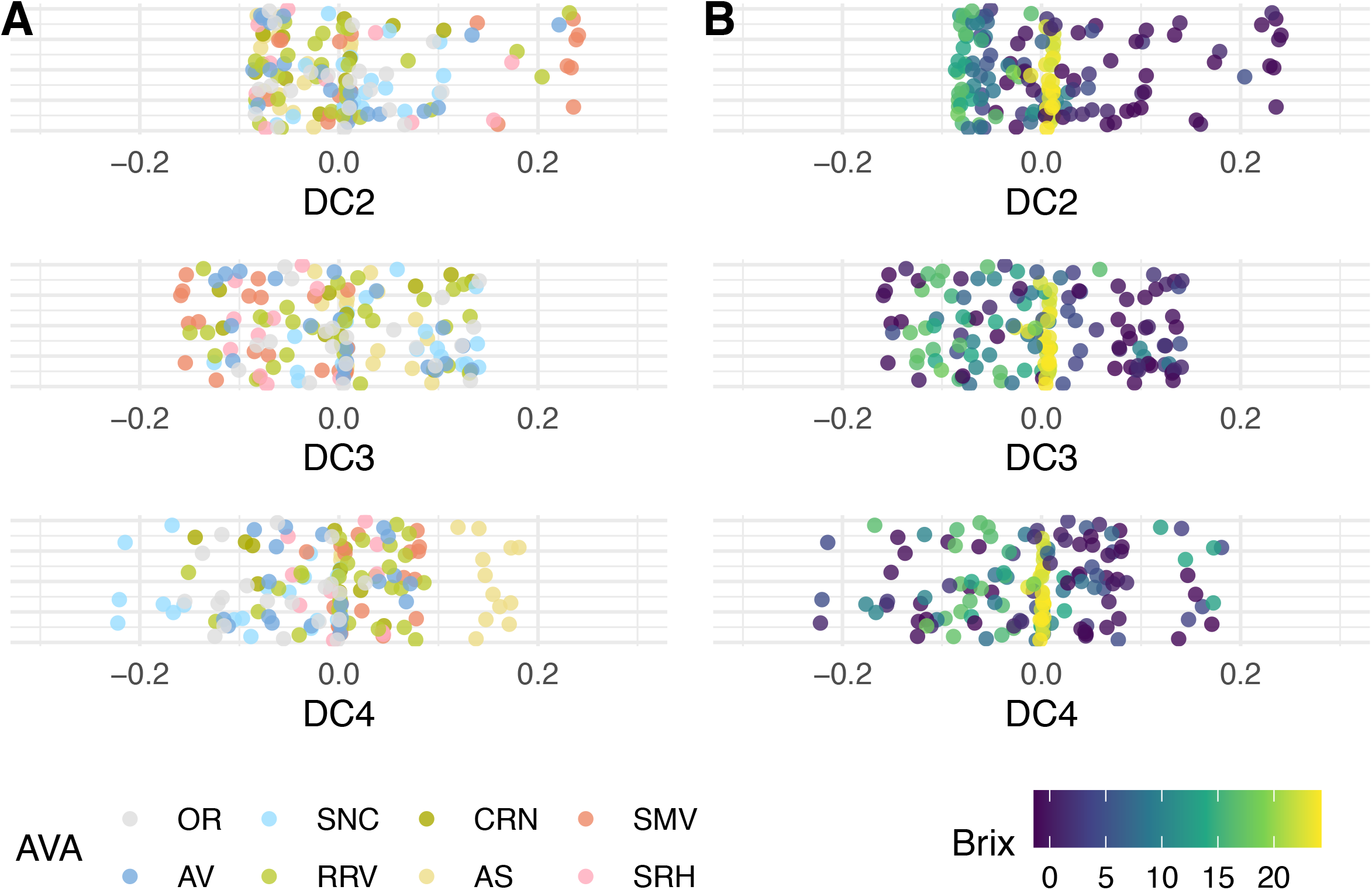
One-dimensional plots of DC2–DC4 for the 2017 vintage. Plots are colored by **A**) American Viticultural Area (AVA) and **B**) Brix. Diffusion components capture different latent structure in the data set. Unlike in the 2019 vintage, samples in lower diffusion components do not cluster, potentially indicating a data quality issue that was remediated in the 2019 vintage using UMI barcodes.

**Figure S3:**
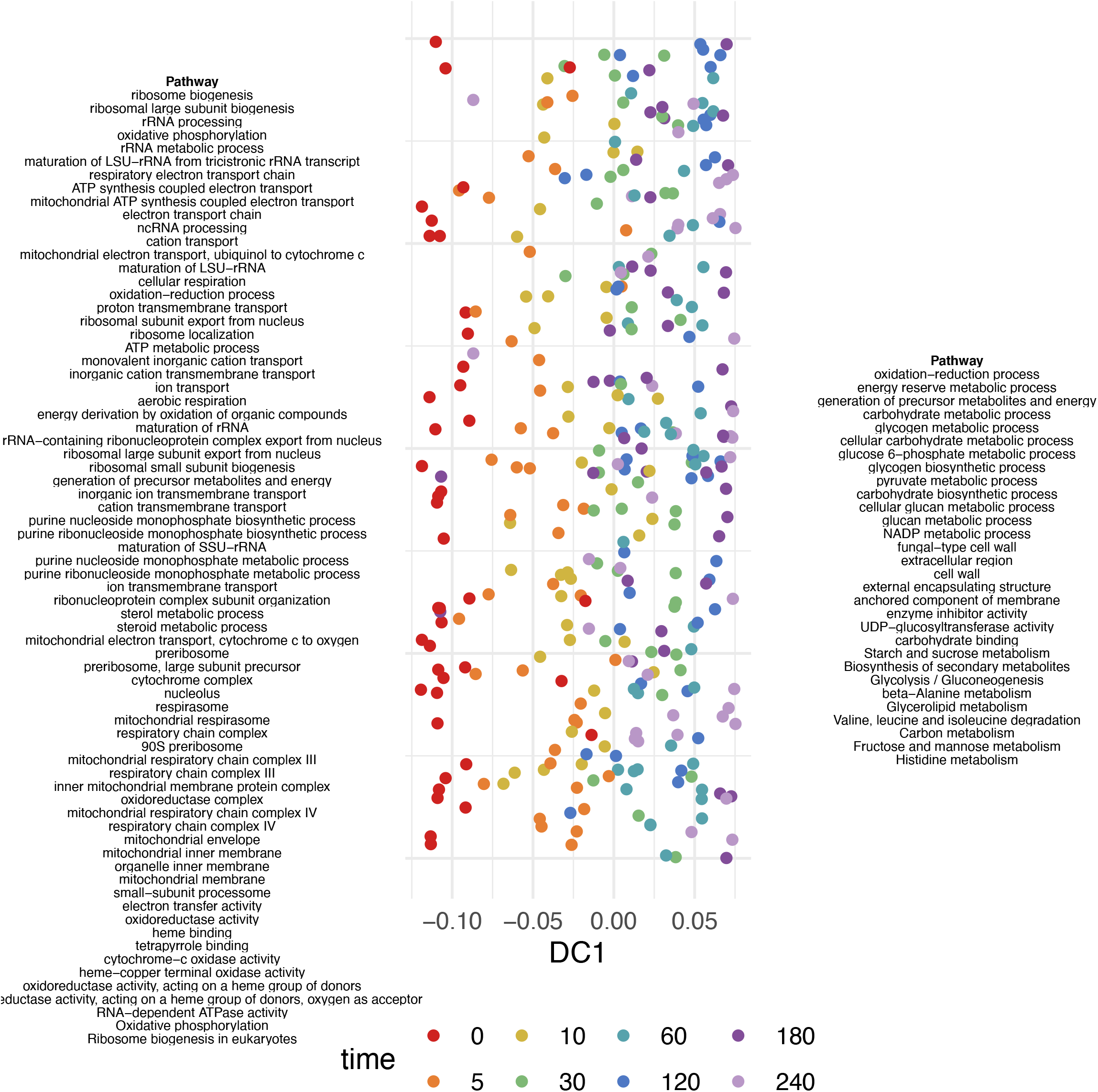
Gene set enrichment for differentially expressed genes along DC1 during onset of hypoxia. All enriched categories with p < 0.05 after Bonferroni correction are shown. Pathways on the right side of the figure are induced in samples with a high diffusion component value, while pathways on the left of the figure are induced in samples with a low diffusion component value.

**Figure S4:**
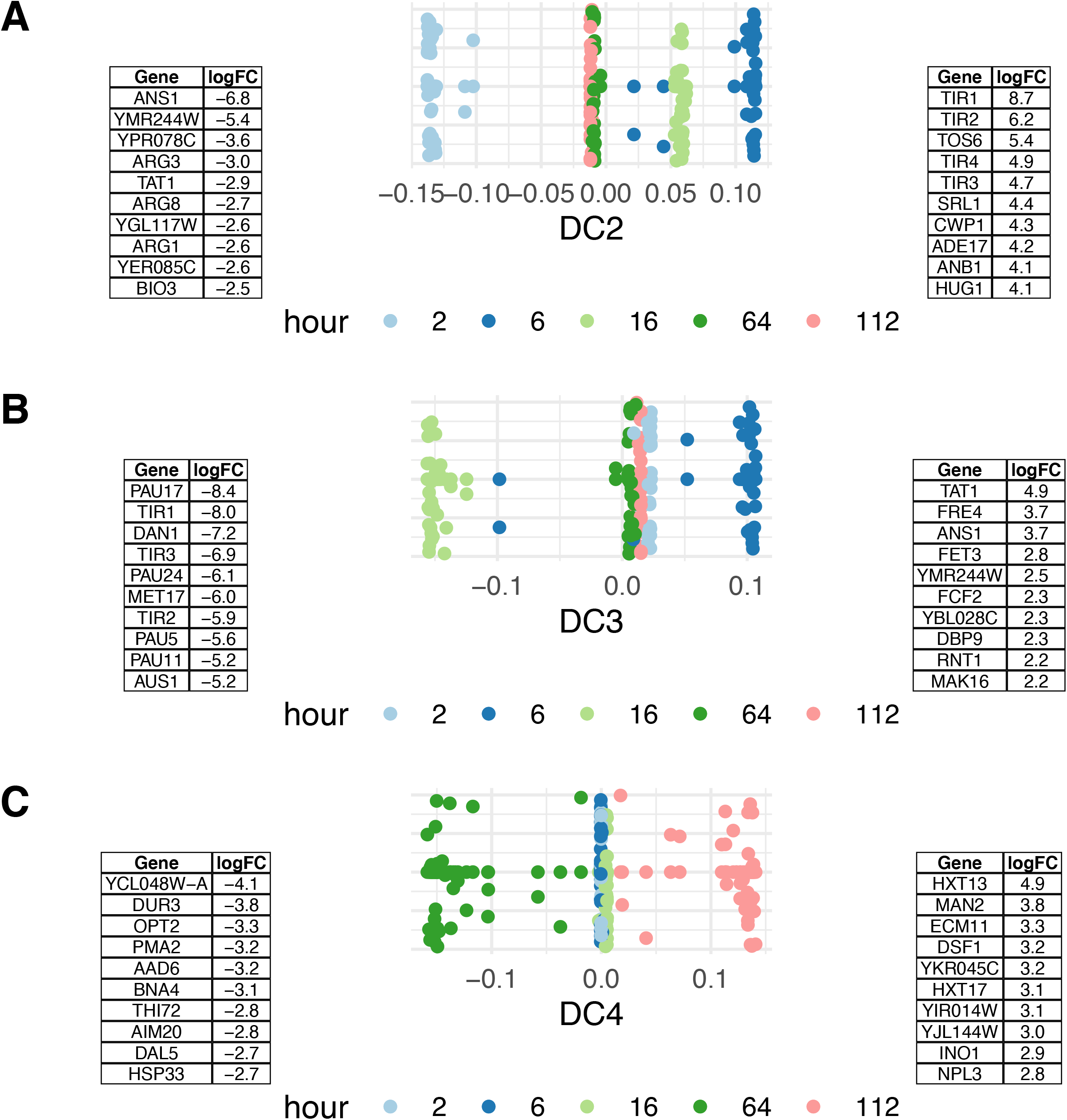
Top 10 genes induced along each diffusion components that separate stages of fermentation in the 2019 vintage. Sites on the diffusion map are colored by hour at which samples were taken. Full gene lists are provided in Supplemental Table 1.

**Figure S5:**
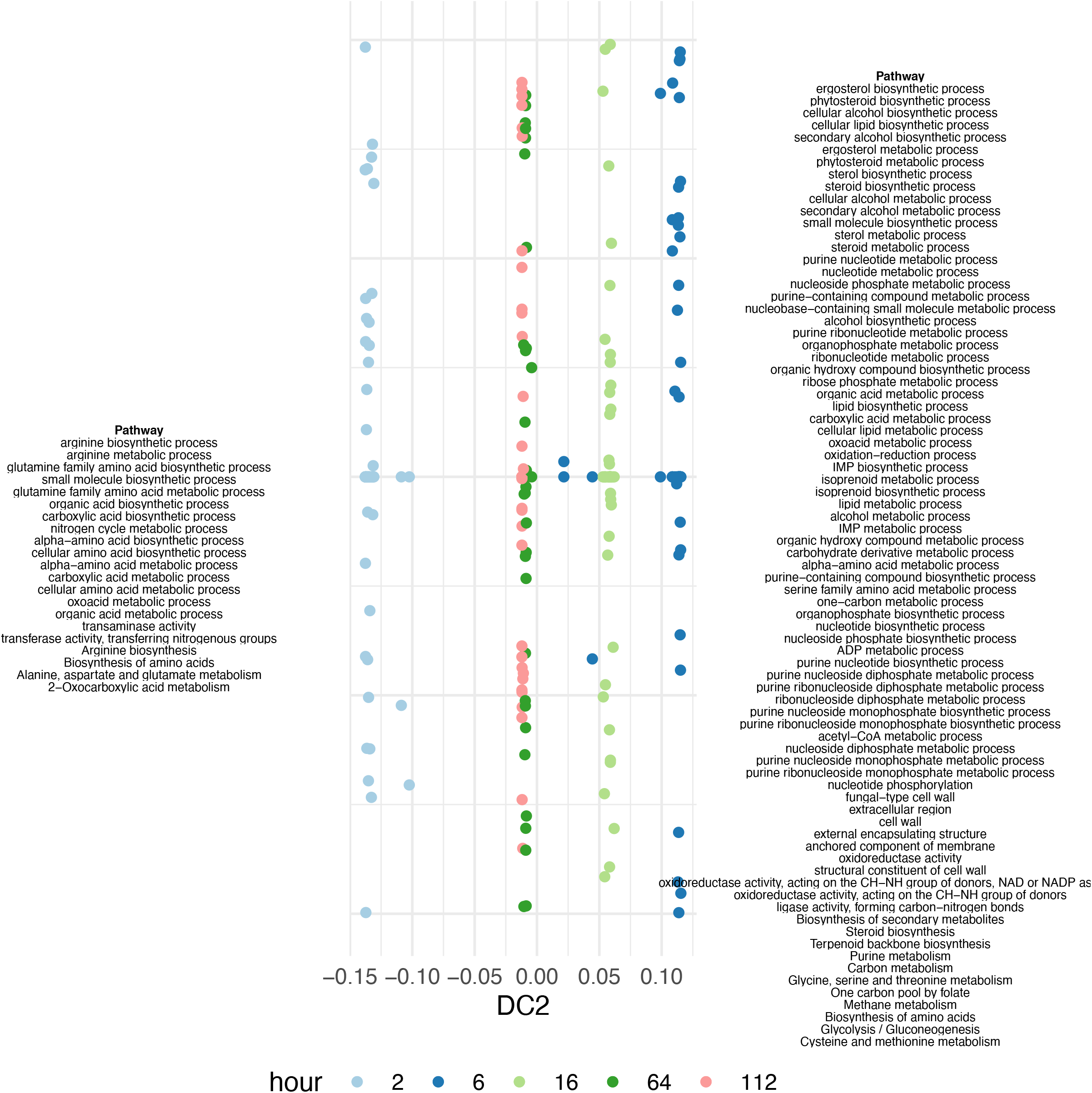
Gene set enrichment for differentially expressed genes along DC2 in the 2019 vintage. All enriched categories with p < 0.05 after Bonferroni correction are shown. Pathways on the right side of the figure are induced in samples with a high diffusion component value, while pathways on the left of the figure are induced in samples with a low diffusion component value.

**Figure S6:**
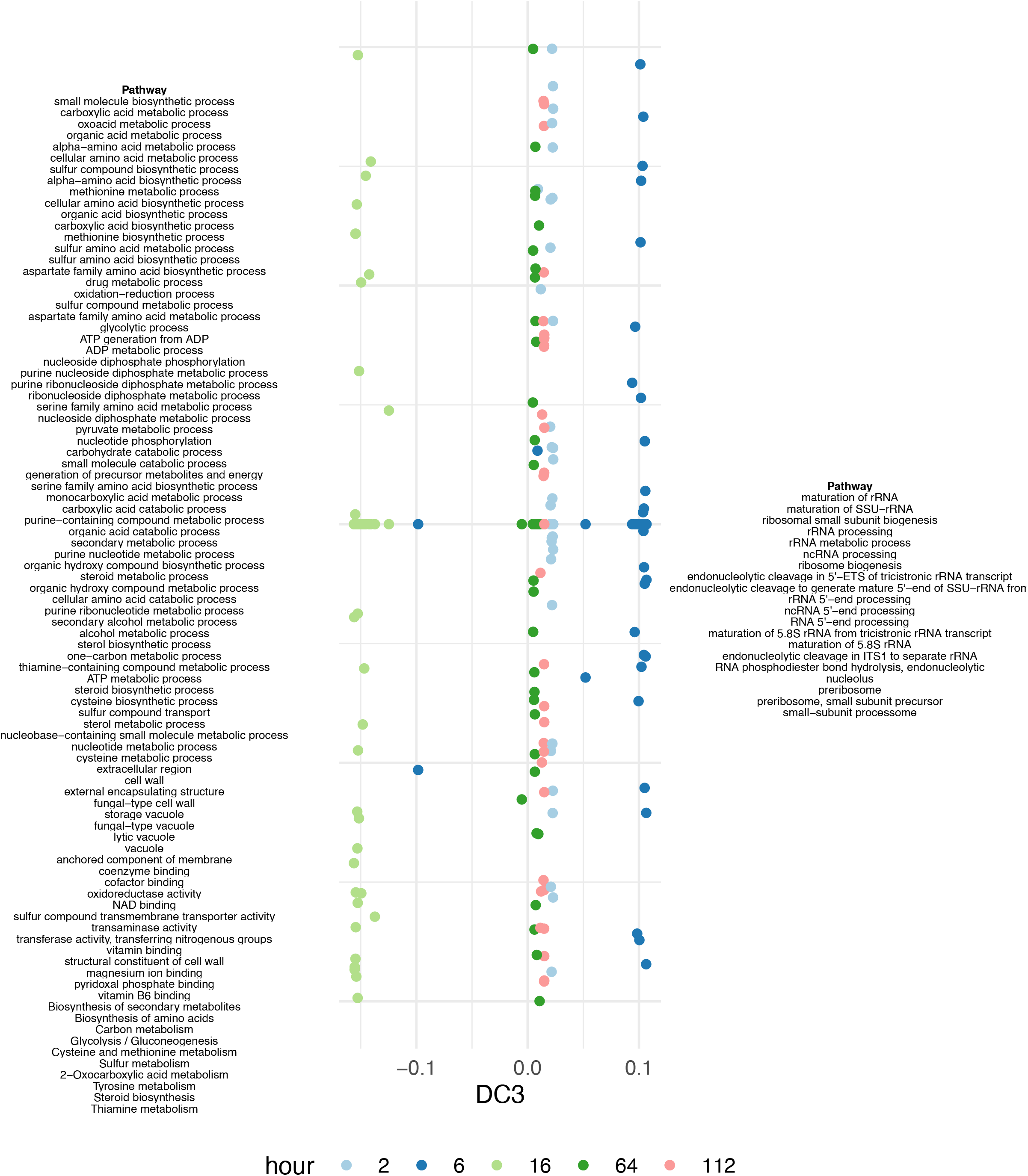
Gene set enrichment for differentially expressed genes along DC3 in the 2019 vintage. All enriched categories with p < 0.05 after Bonferroni correction are shown. Pathways on the right side of the figure are induced in samples with a high diffusion component value, while pathways on the left of the figure are induced in samples with a low diffusion component value.

**Figure S7:**
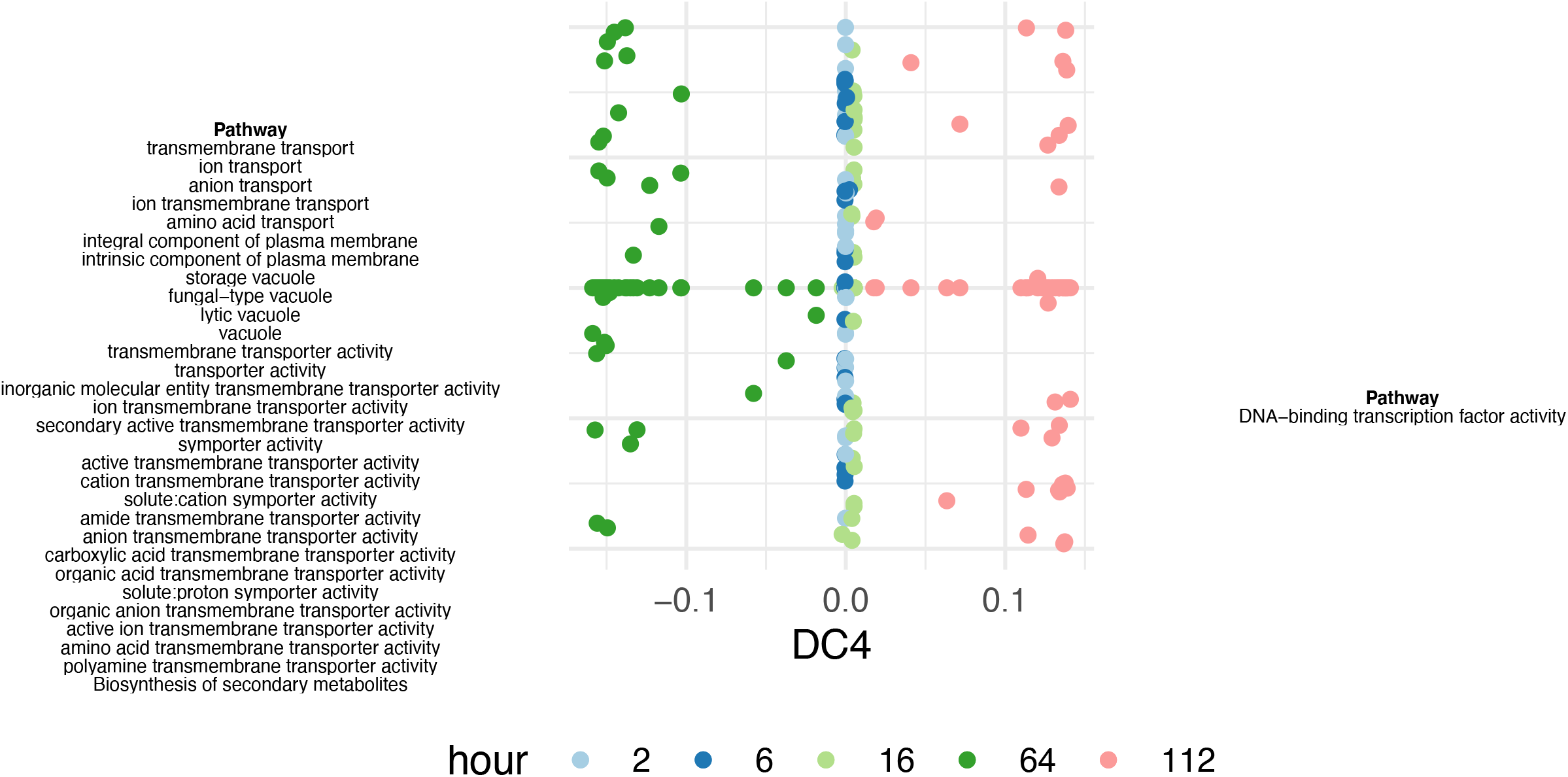
Gene set enrichment for differentially expressed genes along DC4 in the 2019 vintage. All enriched categories with p < 0.05 after Bonferroni correction are shown. Pathways on the right side of the figure are induced in samples with a high diffusion component value, while pathways on the left of the figure are induced in samples with a low diffusion component value.

**Figure S8:**
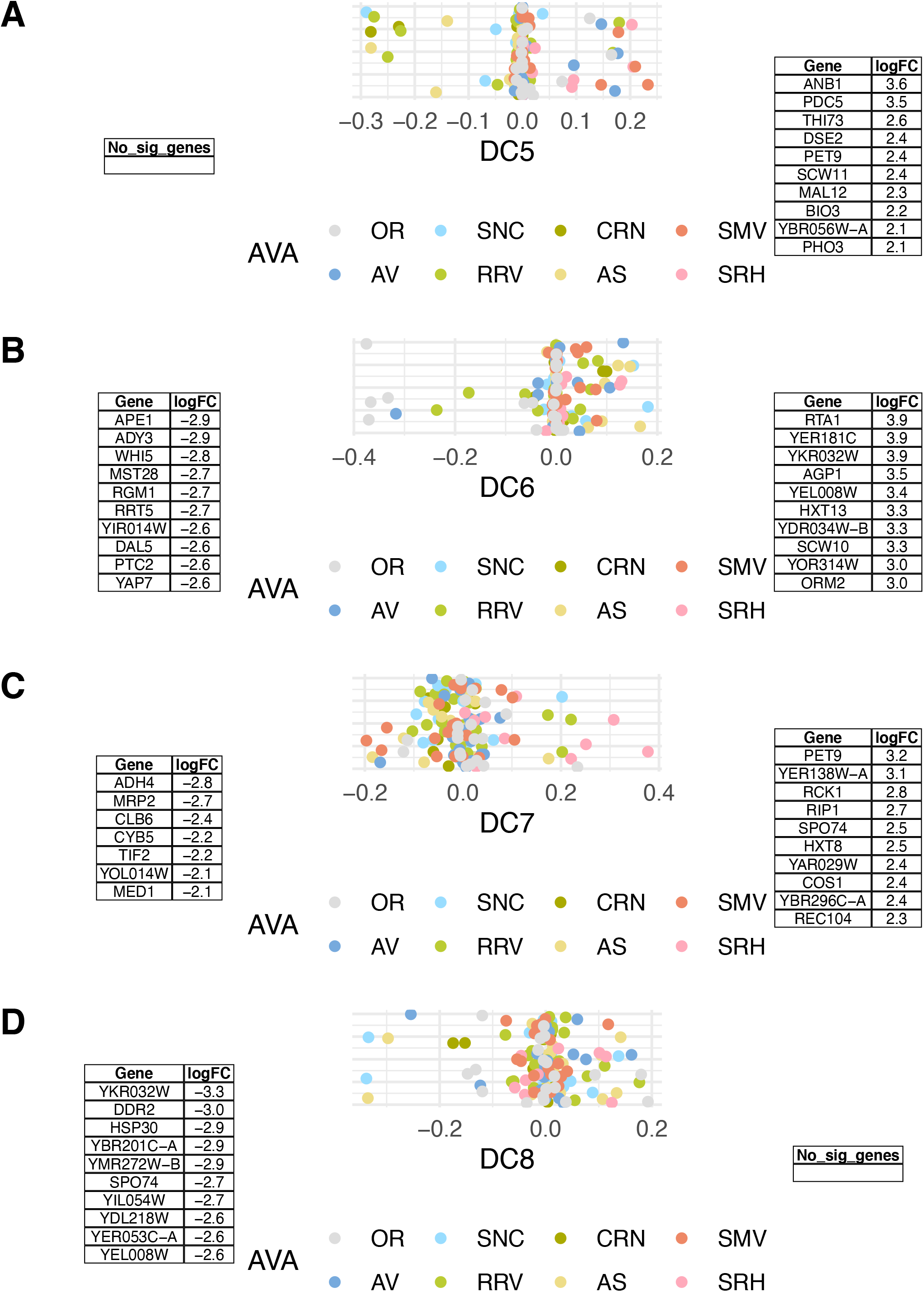
Genes induced along each diffusion components that separate fermentations within a stage of fermentation in the 2019 vintage. Sites on the diffusion map are colored by American Viticultural Area (AVA) of each sample. Full gene lists are provided in Supplemental Table 1.

**Figure S9:**
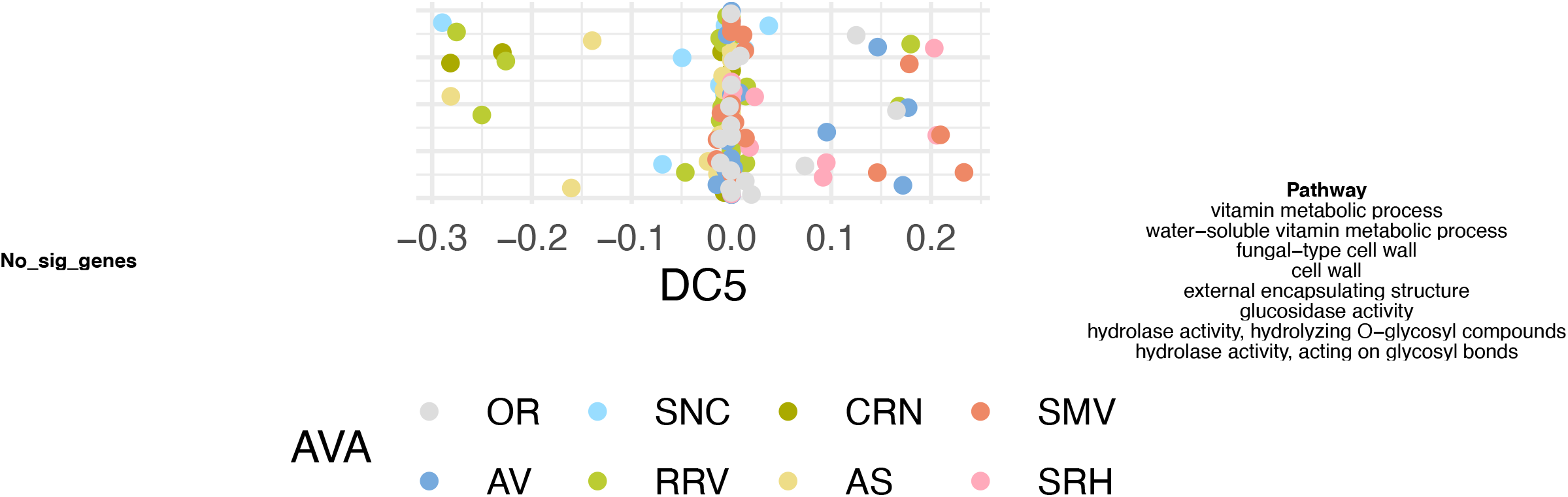
Gene set enrichment for differentially expressed genes along DC5 in the 2019 vintage. All enriched categories with p < 0.05 after Bonferroni correction are shown. Pathways on the right side of the figure are induced in samples with a high diffusion component value, while pathways on the left of the figure are induced in samples with a low diffusion component value.

